# Regulation of neuronal progenitor delamination by dynein-driven post-Golgi apical transport

**DOI:** 10.1101/2021.07.23.453475

**Authors:** J.B. Brault, S. Bardin, M. Lampic, J.A. Carpentieri, L. Coquand, M. Penisson, Hugo Lachuer, G.S. Victoria, S. Baloul, G. Boncompain, S. Miserey-Lenkei, V. Fraisier, F. Francis, F. Perez, B. Goud, A. D. Baffet

## Abstract

Radial glial (RG) cells are the neural stem cells of the developing neocortex. Apical RG (aRG) cells can delaminate to generate basal RG (bRG) cells, a cell type associated with human brain expansion. Here, we report that this delamination is regulated by the post-Golgi secretory pathway. Using *in situ* subcellular live imaging, we show that post-Golgi transport of RAB6+ vesicles occurs toward the minus ends of microtubules and depends on dynein. We demonstrate that the apical determinant Crumbs3 (CRB3) is also transported by dynein. Double knockout of RAB6A/A’ and RAB6B impairs apical localization of CRB3, and induces a retraction of aRG cell apical process, leading to delamination and ectopic division. These defects are phenocopied by knock-out of the dynein activator LIS1. Overall, our results identify a RAB6-dynein-LIS1 complex for Golgi to apical surface transport in aRG cells, and highlights the role of this pathway in the maintenance of neuroepithelial integrity.

## Introduction

In the developing neocortex, all neurons derive from neural stem cells called radial glial (RG) progenitor cells^1, 2^. These highly elongated cells also serve as tracks for the migration of newborn neurons into the cortical plate. Two types of RG cells have been identified: apical RG (aRG) cells (also known as vRG cells), located in the ventricular zone (VZ), and basal RG cells (bRG cells, also known as oRG cells) located in the subventricular zone (SVZ)^3–5^ (Fig. 1a). aRG cells are common to all mammalian species while bRG cells, which originate from aRG cells, are rare in lissencephalic species such as mice but abundant in gyrencephalic species, including humans^6–8^. aRG cells are tightly connected to each other by adherens junctions and form a pseudostratified epithelium lining the ventricle^9^. They are highly polarized and display an apical process extending to the ventricular surface, and a long basal process, connecting to the pial surface (Fig. 1a). Several studies have illustrated that apicobasal polarity is critical for the maintenance of aRG cells, and that its alteration can lead to aRG cell delamination from the neuroepithelium and to the generation of bRG-like cells^10–14^. In ferrets, the cell adhesion molecule cadherin 1 is downregulated at the critical period of bRG cell generation and its knockdown is sufficient to induce bRG cell generation^15^.

**Figure 1.**
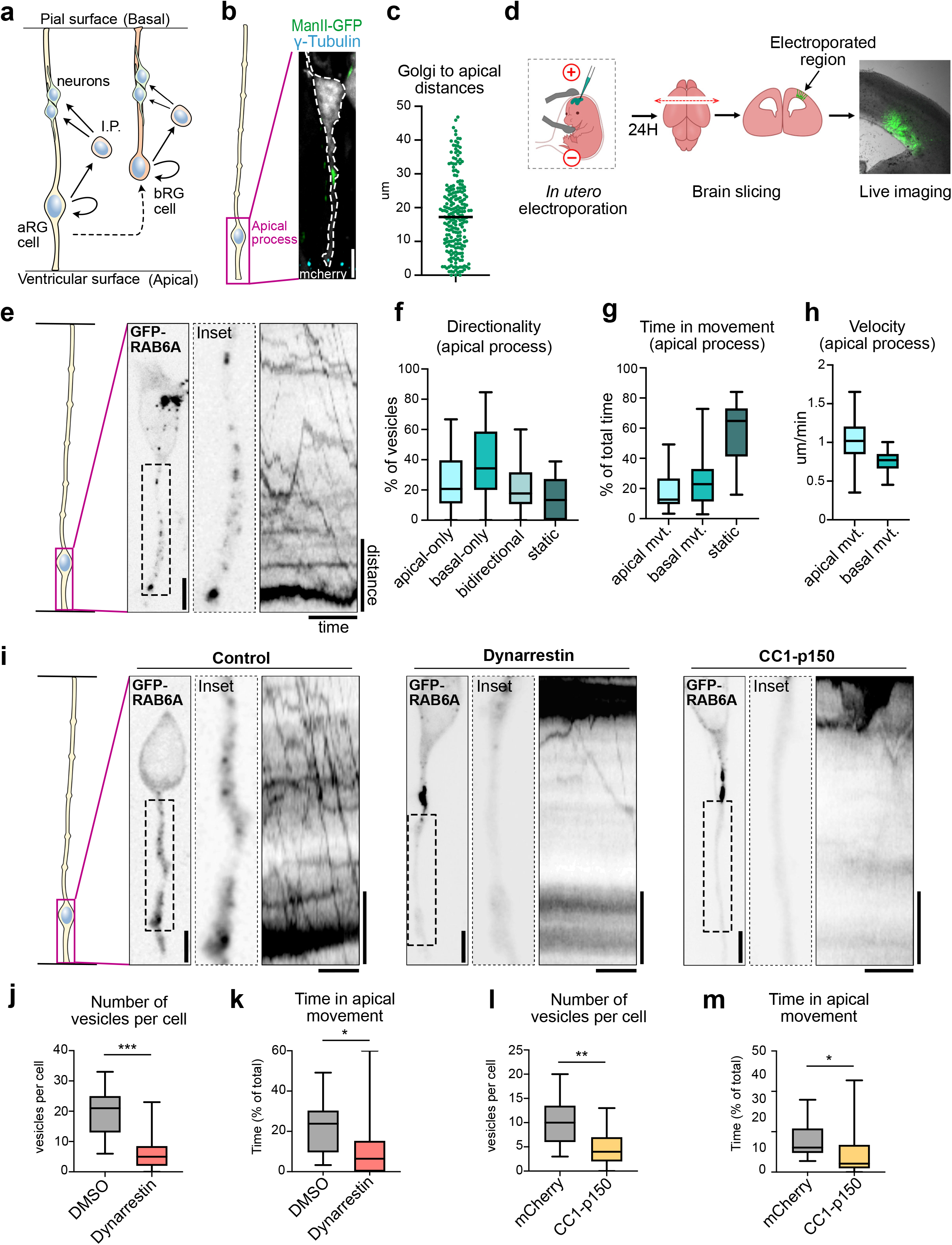
Apical transport of RAB6A+ post-Golgi vesicles is driven by dynein. **a.** Schematic representation of cortical neurogenesis. Apical radial glial (aRG) cells are epithelial cells and the main neuronal progenitors in mouse. Basal radial glial (bRG) cells are rare in mouse but are the most abundant progenitor population in human. They have delaminated from the neuroepithelium. I.P.: Intermediate Progenitor. **b.** Localization of the Golgi apparatus (ManII-GFP) and the centrosome ("-tubulin) in E15.5 mCherry-electroporated radial glial cell. The Golgi apparatus is localized basally, away from the centrosome. Scale bar = 5µm. **c.** Average distance between the apical-most part of the Golgi apparatus and the apical surface in aRG cells. N=224 cells from 3 independent brains. **d.** Schematic representation of *in utero* electroporation and live imaging procedure in the mouse developing cortex. **e.** Live imaging of GFP-RAB6A in aRG cells at E15.5 allows tracking of individual RAB6A+ vesicles *in situ*, from the basal Golgi apparatus towards the apical surface. Scale bar = 5µm. Distance = 5µm, time = 30 seconds. **f.** RAB6A+ vesicle directionality in apical processes of aRG cells over one-minute movies. **g.** Relative time spent by RAB6A+ vesicles in apical, basal or static phases. **h.** Velocity of apically and basally moving RAB6A+ vesicles. (**f, g, h**) N= 388 vesicles from 30 cells. **i.** Live imaging of GFP-RAB6A in control, dynarrestin-treated and CC1-p150-expressing aRG cells at E15.5. Scale bars = 5µm. Distance = 5µm, time = 30 seconds. **j.** Number of RAB6A+ vesicles in the apical process of DMSO and dynarrestin-treated mouse aRG cells. **k.** Relative time spent by RAB6A+ vesicles in apical movement phase, in DMSO and dynarrestin-treated mouse aRG cells. **l.** Number of RAB6A+ vesicles in the apical process of mCherry and CC1-p150-expressing aRG cells. **m.** Relative time spent by RAB6A+ vesicles in apical movement phase, in mCherry and CC1-p150-expressing aRG cells. (**j, k, l, m**) 216 vesicles from N=11 cells for DMSO, 145 vesicles from N=25 cells for dynarrestin, 173 vesicles from N=17 cells for mCherry control, 71 vesicles from N=15 cells for CC1-p150. Mann–Whitney *U* test *p ≤ 0.05, **p ≤ 0.01, *** p ≤ 0.001.

Epithelial polarity is controlled by the PAR, Crumbs and Scribble complexes which mutually interact to generate and maintain apical and basolateral domains. The Crumbs complex is composed of CRB, PALS1, PATJ and is a major apical domain determinant^16^. In the mouse developing neocortex, knock-out of CRB1 and CRB2 leads to an alteration of aRG cells apical junctions, while knockout of PALS1 causes severe polarity defects, apoptotic cell death and microcephaly^17, 18^. The establishment and maintenance of epithelial polarity also rely on polarized trafficking along the biosynthetic/secretory pathway. Newly synthesized transmembrane proteins are sorted in the Golgi apparatus/TGN (*Trans*-Golgi Network) and are routed towards the apical or basolateral domains of epithelial cells, possibly transiting through endosomal compartments^19^. In particular, the secretory pathway is essential for the apical targeting of newly-synthesized CRB, the only transmembrane protein of the apical polarity complex^20^.

RAB6 is a Golgi/TGN-associated small GTPase which controls both anterograde and retrograde transport, from and towards the Golgi apparatus^21^. Three RAB6 paralogs have been identified: ubiquitous RAB6A (and its splicing variant RAB6A’), RAB6B, predominantly expressed in the brain, and RAB6C, encoded by a primate-specific retrogene and involved in cell cycle progression^21, 22^. In non-polarized cells, RAB6A is associated with most - if not all - post-Golgi vesicles, irrespective of the transported cargo, suggesting that RAB6A is a general regulator of post-Golgi trafficking^23^. The exact role of RAB6B is poorly known but evidence exist that it acts redundantly with RAB6A in the secretory pathway^24^. RAB6-positive (RAB6+) secretory vesicles are transported to the cell surface by two plus end-directed kinesins, KIF5B and KIF13B^25^. Retrograde transport towards the Golgi apparatus or the endoplasmic reticulum (ER) is driven by dynein^26, 27^. RAB6 recruits dynein and its partner dynactin through Bicaudal-D (BicD) adaptor proteins, leading to dynein activation and processive movement along microtubules^28–32^. Dynein activity is further regulated by LIS1^33–35^, the dysfunction of which being the most common cause of human lissencephaly^35, 36^. LIS1 activates dynein, but can subsequently be released from an idling complex by RAB6 for processing movement^37^.

In polarized epithelial cells, the machinery controlling trafficking from the Golgi apparatus towards the apical surface is unclear. Conflicting reports have involved both plus-end directed and minus-end directed microtubule motors^38–42^. This is largely due to the limited ability to resolve vesicular transport and post-Golgi trafficking events in polarized epithelial cells, because of the small size of these cells and to the thickness of epithelial tissues. Here, using a method for subcellular live imaging within embryonic brain slices, we show that apical transport of post-Golgi RAB6+ vesicles is driven by dynein. *RAB6A/B* double KO leads to aRG cell delamination during interphase and to the formation of proliferating bRG-like cells. *LIS1* loss of function largely phenocopies *RAB6A/B* dKO, indicating that the RAB6-dynein-LIS1 apical trafficking pathway is required for preventing aRG cell delamination. Finally, we provide evidence that this pathway is critical for the apical transport of the major polarity determinant CRB3 in aRG cells.

## Results

### Post-Golgi apical trafficking occurs towards the minus ends of microtubules

aRG cells are highly elongated cells and undergo interkinetic nuclear migration (INM), a process by which their nuclei translocate basally, before migrating back to the apical surface for mitosis^38, 39^. As a consequence, the average distance between the Golgi apparatus, which follows the nucleus, and the apical surface, where the centrosome is located, is 17.84 !m, ranging from 0 to 46.81 !m, depending on the stage of INM (**Fig. 1b,c**) ^40^. To investigate post-Golgi transport in these cells, we developed an approach for subcellular live imaging within thick organotypic brain slices^41^ (**Fig. 1d**). aRG cells are electroporated with fluorescent reporters *in utero* and, following 24 hours of expression, brains are sliced and mounted for imaging on a CSU-W1 spinning disk microscope equipped with a high working distance 100X objective (see methods). This approach allowed the visualization of growing microtubule plus ends in cells expressing the plus end tracking protein EB3. We confirmed our previous results, i.e. the unipolar organization of the microtubule network with over 99% of plus ends growing in the basal direction, from the pericentrosomal apical surface^41^ (**Extended Data Fig. 1a**, **Supplemental Video 1**). Notably, virtually no microtubules emanating from the Golgi area were observed to grow apically.

To visualize post-Golgi transport vesicles, we electroporated aRG cells *in utero* with a GFP-RAB6A expressing plasmid. The construct was expressed at low levels to avoid cytosolic accumulation, and 3-5 planes were imaged to capture the entire apical process, leading to a temporal resolution of 600-1000 ms. GFP-RAB6A marked the Golgi apparatus, which sometimes appears fragmented as previously reported in these cells^40^, as well as small and dynamic vesicular structures that could often be observed budding from the Golgi (**Extended data Fig. 1b, Supplemental Video 2**). Live imaging within the apical process (between the Golgi and the apical surface) revealed that RAB6A+ vesicles were bidirectional (**Fig. 1e, Supplemental Video 3**). Highly dynamic RAB6A+ vesicles could also be observed within the basal process (above the nucleus), where they also appeared highly dynamic (**Extended data Fig. 1c, Supplemental Video 4**). In the apical process, manual tracking of individual RAB6A+ vesicles revealed that, throughout one-minute movies, 39% displayed basal movement (towards the Golgi apparatus), 25% apical movement (towards the apical surface), 21% bidirectional movement and 15% were static (**Fig. 1f**). These vesicles spent 24% of their time moving in the basal direction, 18% moving in the apical direction, and 58% not moving (**Fig. 1g**). Apically-moving RAB6A+ vesicles moved faster than basally-moving ones, in agreement with faster minus-end transport reported in non-polarized cells^25, 42^ (**Fig. 1h**). Including pauses, RAB6A+ vesicles traveled on average 32.3 !m per minute. They were often observed to disappear at the apical surface, suggesting apical fusion events, either with the plasma membrane or with another compartment (**Extended data Fig. 1d, Supplemental Video 5**). Together, these results reveal that RAB6A+ vesicles traffic in a highly bidirectional manner between the perinuclear Golgi apparatus and the apical surface, which they reach following transport directed towards microtubule minus ends.

### Apical transport of post-Golgi RAB6A+ vesicles is driven by dynein

We next asked whether post-Golgi apical transport of RAB6A+ vesicles was dependent on the minus end microtubule motor dynein. To test this, we treated brain slices with the dynein inhibitor dynarrestin, prior to live imaging^43^. Because of its short stability, a new batch of dynarrestin was dissolved prior to each experiment, and validated in parallel for Golgi dispersal in RPE-1 cells (**Extended data, Fig. 1e**). Dynarrestin treatment in aRG cells led to a drastic inhibition of the trafficking of RAB6A+ vesicles into the apical process, as compared to DMSO-treated cells (**Fig. 1i**, **Supplemental Videos 6, 7**). The total number of RAB6A+ vesicles observed within the apical process was severely reduced (**Fig. 1j**). This result suggests that, in the absence of dynein activity, the balance between opposing motors was shifted towards kinesin-dependent transport in the basal direction, leading to an emptying of the apical process. The vesicles that did manage to enter the apical process performed substantially less apically-directed movements (**Fig. 1k**). On the contrary, RAB6A+ vesicles in the cell soma and in the basal process remained highly mobile.

To confirm these results, we next overexpressed a truncated form of p150*^Glued^* (CC1-p150), which acts as a dominant-negative for the dynactin complex^44^. Expression of CC1-p150 for 24 hours in aRG cells led to a very similar outcome, impairing the localization of RAB6A+ vesicles into the apical process (**Fig. 1i, l Supplemental Video 8**). As for dynarrestin treatment, apical movement of RAB6A+ vesicles located within the apical process was markedly reduced (**Fig. 1m**). Mobile RAB6A+ vesicles were still abundant in the soma and basal process. In both cells treated with dynarrestin or overexpressing CC1-p150, the speed of RAB6A+ vesicles that were still moving was unaltered within the apical process (**Extended data, Fig. 1f, g**). Together, these results indicate that post-Golgi RAB6A+ vesicles travel towards the apical surface of aRG cells along a uniformly polarized microtubule network *via* dynein-based transport.

### RAB6A/B double knockout causes microcephaly

We next investigated the consequences of RAB6 loss of function on neocortical development, neuroepithelial integrity, and apical cargo delivery. We confirmed RAB6A/A’ and B expression in the developing brain, and observed that RAB6B expression strongly rises from E11.5, while RAB6A/A’ levels remain constant (**Extended data, Fig. 2a**). Because constitutive knock-out of *RAB6A* (coding for the two isoforms RAB6A and RAB6A’) leads to early developmental lethality^45^, we previously generated a Cre-inducible KO mouse model^46^. Dorsal cortex-specific depletion of *RAB6A*, using the *Emx1-Cre* driver, did not lead to any observable phenotype on neocortex development. To test for redundancy, we therefore generated a constitutive KO mouse for *RAB6B*, using Crispr-Cas9. We obtained two lines, a 279 bp inversion affecting in exons 3 and 4, and a 1 bp deletion in exon 2, both leading to a premature stop codon. Both lines were viable and, as for conditional *RAB6A* KO, did not display any observable alterations of neocortex development or aRG cell polarity. We therefore generated *RAB6A/B* double KO (*RAB6A/B* dKO) animals. Efficient protein depletion for RAB6A/A’ and RAB6B in the embryonic cortex was verified by western blot (**Fig. 2a**), residual RAB6A/A’ signal in *RAB6A/B* dKO mice being likely due to the presence of non-Cre expressing cells in the protein extract. Strikingly, *RAB6A/B* dKO mice were severely microcephalic. At P0, the cortical area as well as the cortical thickness of double mutant animals were reduced by half, while single KOs were unaffected (**Fig. 2b, c, d, e**). Reduced brain size was likely the consequence of increased levels of apoptotic cell death observed in *RAB6A/B* dKO (**Fig. 2f**). Neuronal positioning was also strongly affected, with layer II-III neurons (CDP+) dispersed throughout the neocortex, suggesting impaired neuronal migration (**Fig. 2d**). Therefore, loss of RAB6A/A’ and RAB6B leads to microcephaly and altered neuronal positioning.

**Figure 2.**
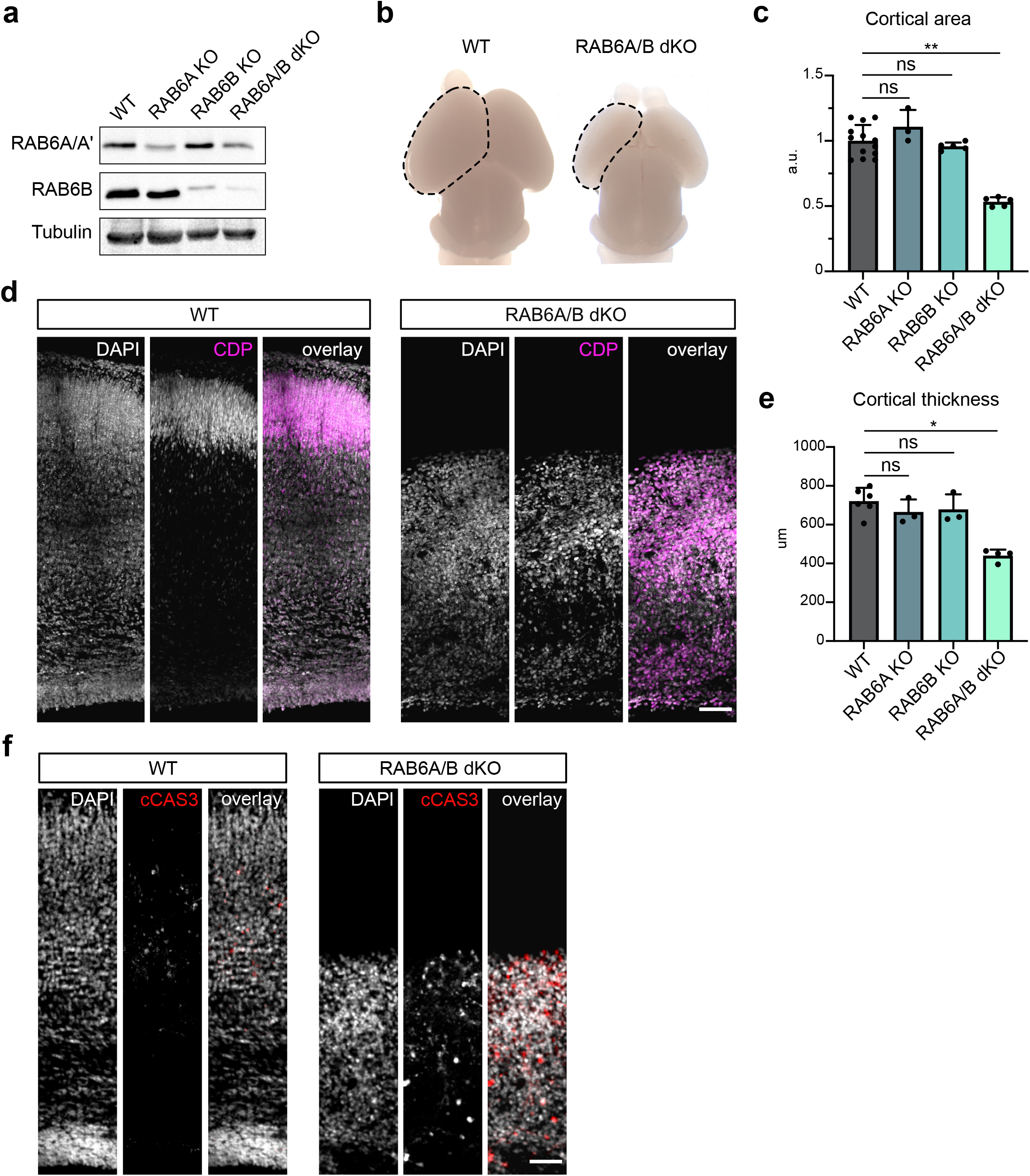
*RAB6A/B* double knockout causes microcephaly. **a.** Western blot analysis of RAB6A/A’ and RAB6B protein levels in *WT*, *Emx1-Cre; RAB6^loxP/loxP^* (*RAB6A* KO), *RAB6B^-/-^* (*RAB6B* KO) and *Emx1-Cre; RAB6A^loxP/loxP^*; *RAB6B^-/-^* (*RAB6A/B* dKO) E15.5 cortical extracts. **b.** P0 WT and *RAB6A/B* dKO brains. A cortical hemisphere is circled (dotted lines). **c.** Cortical area in WT (N=13), *RAB6A* KO (N=2), *RAB6B* KO (N=5) and *RAB6A/B* dKO (N=5) at P0. **d.** WT and *RAB6A/B* dKO brains stained for layer II/III marker CDP at P0. Scale bar = 100µm. **e.** Cortical thickness (μm) in WT (N=6), *RAB6A* KO (N=3), *RAB6B* KO (N=3) and *RAB6A/B* dKO (N=4) at P0. **f.** Immunostaining for cleaved Caspase 3 (cCAS3) in WT and *RAB6A/B* dKO brains at P0. Scale bar = 50µm. (**c, e**) Kruskal-Wallis test with a Dunn *post-hoc* test and Benjamini-Hochberg procedure.

### RAB6A/B dKO leads to aRG cell delamination during interphase

We next analyzed *RAB6A/B* dKO embryos to test for neuroepithelial organization defects. To monitor aRG cell positioning defects, we first analyzed the localization of PAX6+ aRG cells. In E15.5 control as well as in single *RAB6A* and *RAB6B* KO brains, aRG cells were concentrated within the VZ. In *RAB6*A/B dKO however, numerous RG cells could be observed above the VZ, suggesting delamination from the neuroepithelium (**Fig. 3a, b**). Moreover, the size of the PAX6+ VZ was reduced, even when normalized to total cortical thickness, further indicating a loss of ventricular aRG cells (**Extended data, Fig. 2b**). The presence of ectopic RG cells was confirmed by the strong increase in the fraction of mitotic RG cells located above the ventricular surface, positive for phospho-Vimentin (p-VIM), which specifically marks mitotic RG cells (**Fig. 3c, d**). This staining further revealed that basally located RG cells had lost their apical process and had therefore detached form the neuroepithelium. They also appeared to have retracted their basal process, but continued to divide. Quantification of the mitotic index of PAX6+ RG cells indeed indicated that ectopic *RAB6A/B* dKO RG cells proliferated at a normal rate (**Fig. 3e**).

**Figure 3.**
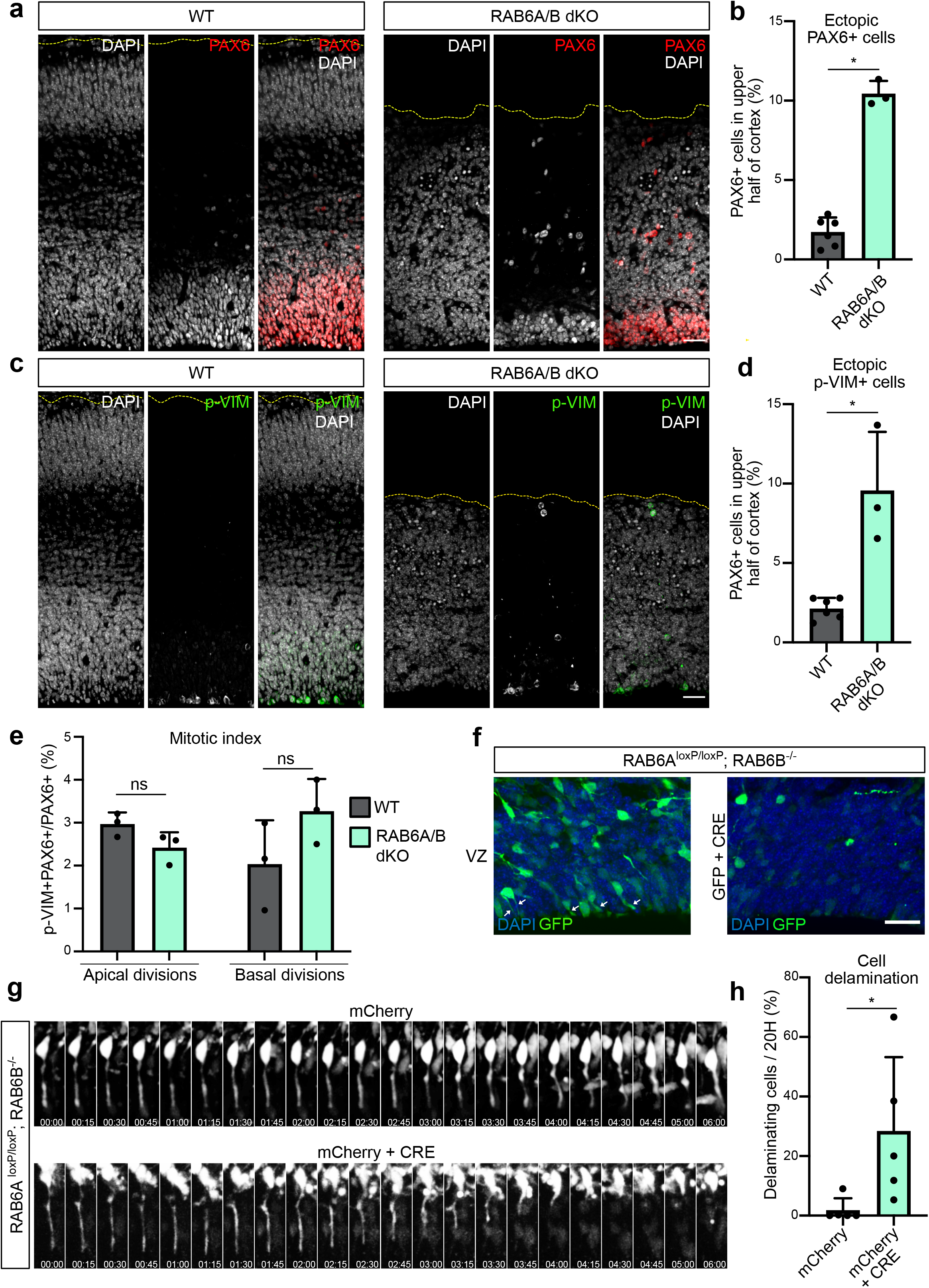
*RAB6A/B* dKO leads to aRG cell delamination during interphase. **a.** PAX6 staining in WT and *RAB6A/B* dKO E15.5 brains. Scale bar = 50µm. **b.** Percentage of PAX6+ cells located in the upper half of the cortex of WT and *RAB6A/B* dKO E15.5 brains. WT: 4282 cells from N=6 brains. *RAB6A/B* dKO: 1241 cells from N=3 brains. Mann–Whitney *U* test. **c.** Phospho-Vimentin (p-Vim) staining in WT and *RAB6A/B* dKO E15.5 brains. Scale bar = 50µm. **d.** Percentage p-VIM+ cells dividing ectopically, in the upper half of the cortex of WT and *RAB6A/B dKO* E15.5 brains. WT: 1713 cells from N=6 brains. *RAB6A/B* dKO: 506 cells from N=3 brains. Mann–Whitney *U* test. **e.** Mitotic index (p-VIM+ PAX6+ / PAX6+ cells) of RG cells dividing apically (at the ventricular surface) or basally (upper half) in WT and *RAB6A/B* dKO E15.5 brains. 3 to 6 brains were analyzed per condition. Apical divisions: N= 1886 cells for WT and 1511 cells for *RAB6A*/*B* dKO. Basal divisions: N=643 cells for WT and 809 cells for *RAB6A/B* dKO. Mann–Whitney *U* test. **f.** Electroporation of *RAB6A^loxP/loxP^*; *RAB6B^-/-^* E14.5 embryos with GFP (control) or GFP + CRE (*RAB6A/B* dKO) and fixation at E18.5. Localization of GFP+ cells in the ventricular zone. White arrows indicate apical processes. Scale bar = 25µm. **g.** Electroporation of *RAB6A^loxP/loxP^*; *RAB6B^-/-^* E14.5 embryos with mCherry (control) or mCherry + CRE (*RAB6A/B* dKO) and live imaging of delamination events at E17.5. **h.** Apical endfoot detachment and retraction events during 20 hours movies in mCherry or mCherry + CRE electroporated cells at E17.5. mCherry: N=46 cells from 5 movies. mCherry + CRE: N=72 cells from 5 movies. Fisher’s exact test, *p ≤ 0.05.

To investigate further whether these ectopic aRG cells had indeed delaminated from the neuroepithelium, we used *in utero* electroporation, which specifically targets the aRG cells and therefore allows to assess the position of these cells and their progeny over time. We electroporated a plasmid coding for the Cre recombinase, as well as GFP, into E14.5 *RAB6A^loxP/loxP^; RAB6B^-/-^* brains, in order to deplete both RAB6A/A’ and B specifically in the GFP-expressing electroporated aRG cells. After 4 days in control GFP-electroporated brains, numerous aRG cells could be observed connected to the ventricular surface by their apical processes. In Cre-expressing brains however, these cells were largely lost, suggesting that they had detached from the neuroepithelium (**Fig. 3f**). To confirm that the presence of basally-localized RG cells was indeed a consequence of apical process detachment during interphase, we live imaged aRG cells 3 days after Cre expression-induced *RAB6A/B* dKO. While the majority of control cells maintained an apical attachment throughout 20 hour-long movies, a high proportion of Cre-expressing *RAB6A/B* dKO RG cells were observed to detach from the neuroepithelium and retract their apical process towards the cell soma (**Fig. 3g, h, Supplemental Videos 9, 10**). Together, these results indicate that double depletion of RAB6A/A’ and B leads to the delamination of RG cells during interphase. These cells however maintain an RG fate and continue to proliferate above the VZ.

### LIS1 knock-out leads to ectopically dividing progenitors

We next asked whether the delamination observed in *RAB6A/B* dKO was a consequence of altered trafficking towards the apical surface, and therefore if altered dynein activity would lead to a similar outcome. To test this, we inactivated *LIS1* in the mouse neocortex, using an inducible KO mouse model^47^. *Emx1-Cre; LIS1^loxP/loxP^* (*LIS1* KO) were severely microcephalic, as previously described^47^. PAX6+ cells in E12.5 *LIS1* KO were found dispersed throughout the entire tissue (**Fig. 4a**). The majority of mitotic RG cells (PAX6+ p-H3+) were localized basally, away from the apical surface where they are normally found, suggesting that they had delaminated (**Fig. 4a, c**). This result was confirmed following p-VIM staining (**Fig. 4b, d**). Therefore, inhibition of dynein through LIS1 loss of function largely phenocopies *RAB6A/B* dKO, suggesting that the RAB6-dynein-LIS1 apical trafficking pathway is required to prevent aRG cell delamination.

**Figure 4.**
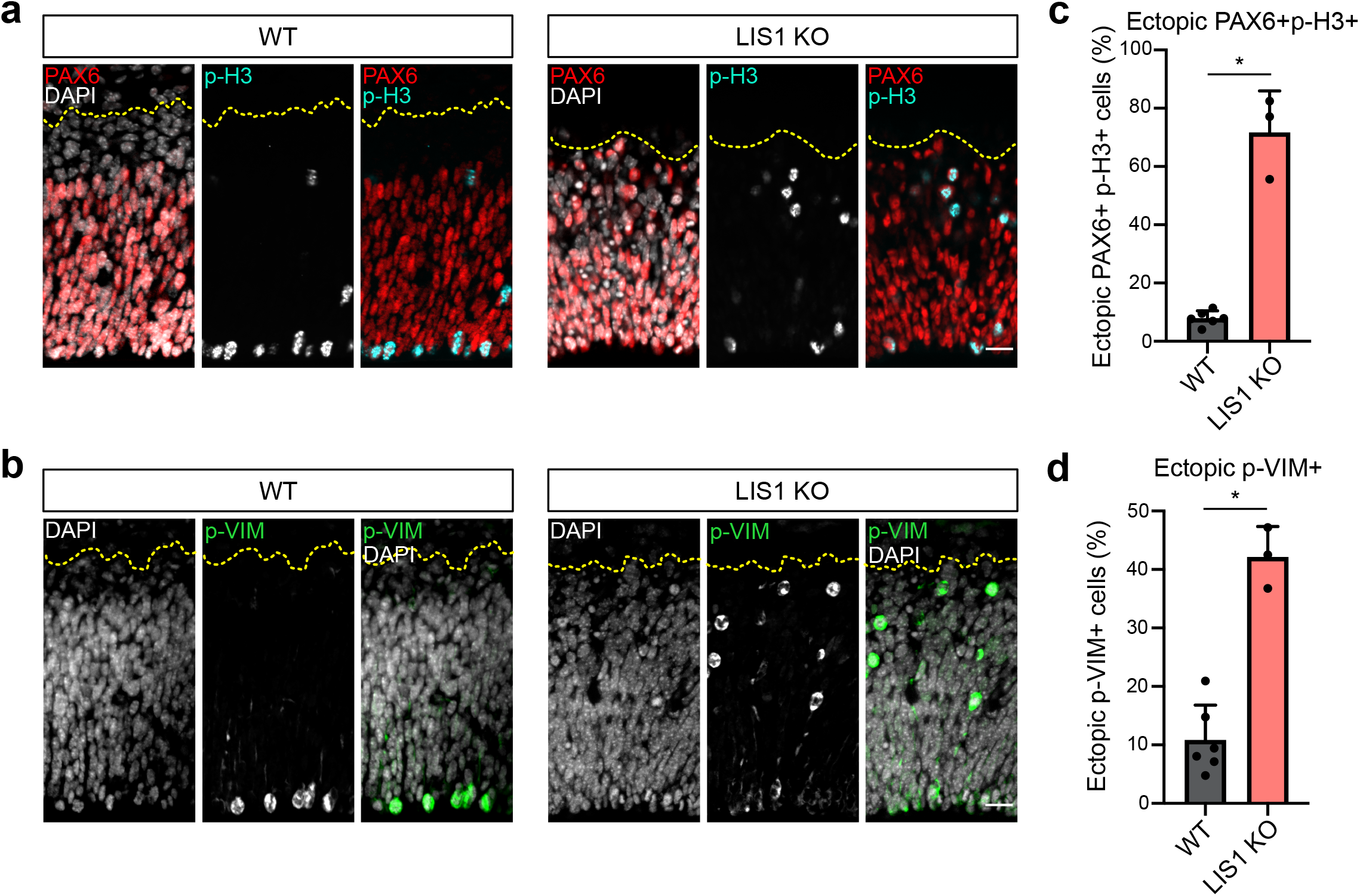
*LIS1* knock-out leads to ectopically dividing progenitors. **a.** PAX6 and phospho-Histone 3 (p-H3) staining in WT and *LIS1* KO E12.5 brains. Cortices were subdivided into 5 bins of equal size along the radial axis. Scale bar = 25µm. **b.** Phospho-Vimentin (p-VIM) staining in WT and *LIS1* KO E12.5 brains. Scale bar = 25µm. **c.** Percentage of PH3+/PAX6+ cells located above the ventricular surface of WT and *LIS1* KO E12.5 brains. WT: 1192 cells from N=6 brains. *LIS1* KO: 589 cells from N=3 brains. **d.** Percentage p-VIM+ cells dividing ectopically, away from the ventricular surface of WT and *LIS1* KO E12.5 brains. WT: 1056 cells from N=6 brains. *LIS1* KO: 879 cells from N=3 brains. (**c, d**) Mann–Whitney *U* test.

### Post-Golgi apical transport of Crumbs is driven by dynein

Interphasic delamination is a consequence of destabilization of the adherens junctions, which are themselves dependent on properly established epithelial polarity. The transmembrane protein CRB is a major determinant of epithelial apical domain polarity and the only one to be transported along the secretory pathway. Accordingly, CRB3, the major Crumbs isoform expressed in mammalian epithelial cells^48^, and its partner PALS1 localize to the apical surface of aRG cells (**Fig. 5a**). We therefore asked whether the RAB6-dynein-LIS1 pathway controls the apical transport of CRB3 in these cells. To distinguish between different trafficking pools –secretory and endolysosomal– we analyzed CRB3 trafficking using the RUSH system^49^ (**Fig. 5b**). This assay allows for the retention of a cargo of interest in the ER and, upon addition of biotin, its release for trafficking along its secretory route. Following *in utero* electroporation, SBP-CRB3-GFP was efficiently retained *in vivo* within the ER and absent from the apical surface of aRG cells, indicating that endogenous biotin levels in mouse were not sufficient to induce its release (**Fig. 5c, d**). To monitor SBP-CRB3-GFP trafficking, brain slices were incubated in the presence of biotin and fixed at different time points. At 20 minutes, CRB3 had arrived at the Golgi apparatus in most aRG cells (95.7±5.2%), and by 60 minutes it strongly accumulated at the apical surface of over 90% of the cells (**Fig. 5c, d**). In one third of the cells, CRB3 was only detected at the apical surface, indicating that most of the protein pool had reached its final location (**Fig. 5c, e**).

**Figure 5.**
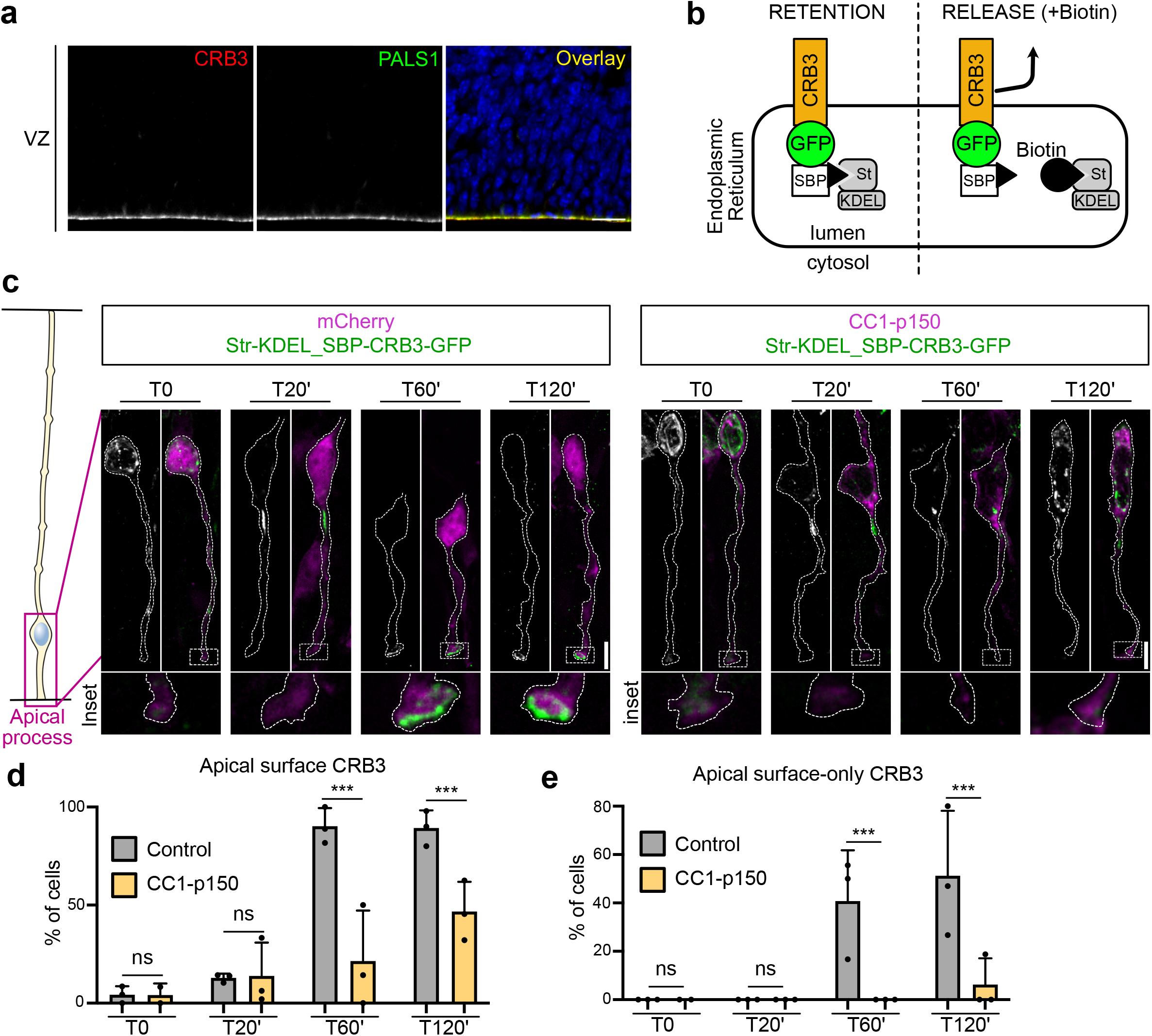
Post-Golgi apical transport of Crumbs is driven by dynein. **a.** Immuno-staining for Crumbs3 (CRB3) and PALS1 in E15.5 embryonic cortex. Scale bar = 25µm. **b.** Schematic representation of the RUSH system. CRB3 is retained in the endoplasmic reticulum until the addition of biotin, which releases it for trafficking. SBP: Streptavidin-binding protein. St: Streptavidin. **c.** RUSH assay for CRB3-GFP in control (mcherry) and dynactin-inhibited radial glial cells (CC1-p150-dsRed), electroporated at E.15.5 and imaged at E16.5. Scale bar = 5µm. **d.** CRB3 localization at the apical surface upon release. **e.** Percentage of cells with 100% of CRB3 signal at the apical surface upon release. (**d, e**) mCherry (control): N= 361 cells. CC1-p150: N=268 cells. Fisher’s exact test and Benjamini-Hochberg procedure, *** p ≤ 0.001.

To test whether post-Golgi transport of CRB3 towards the apical surface relies on dynein, we monitored SBP-CRB3-GFP trafficking in aRG cells expressing the CC1-p150 dominant negative construct. As in control, 20 minutes after biotin treatment, CRB3 reached the Golgi apparatus (in 94.1±3.6% of the cells), but at 60 minutes its trafficking towards the apical surface was severely affected (**Fig. 5c, d**). By 120 minutes, it started to reach the apical surface, although exhibiting a 2-fold decrease compared to control cells. Moreover, almost no CC1-expressing cell showed a localization of the total CRB3 pool at the apical surface, even after 120 minutes, as compared to a third of control cells (**Fig. 5c, e**). Therefore, post-Golgi transport of Crumbs towards the apical surface of aRG cells is driven by the dynein-dynactin complex.

We and others have previously shown that post-Golgi RAB6A+ vesicles contain a wide variety of cargoes^23, 50, 51^. We confirmed here that RAB6A+ vesicles also transport CRB3. HeLa cells expressing CRB3 in the RUSH system were imaged 30 minutes after biotin addition, when CRB3 has reached the Golgi apparatus and begun to exit it. At this timepoint, almost 80% of vesicles containing SBP-CRB3-GFP were positive for mcherry-RAB6A, indicating that CRB3 largely exits the Golgi apparatus within RAB6A+ vesicles (**Extended data, Fig. 3a, b**).

### Apical localization of Crumbs in aRG cells depends of RAB6A/B and LIS1

Finally, we tested the consequence *of LIS1 and RAB6A/B* KO on the steady-state levels of the Crumbs complex at the apical surface of aRG cells. *LIS1* KO brains revealed altered apical localization of CRB3 and its partner PALS1 (**Fig. 6a**). The CRB3 apical signal intensity was reduced, which we quantified using line scan fluorescent intensity measurements (**Fig. 6b**). Moreover, we observed the frequent appearance of patches that were completely devoid of CRB3 and PALS1. We quantified the number of empty patches along the ventricular surface, which were completely absent in control embryos but occurred at a frequency of 8.2 per mm in *LIS1* KO embryos (**Fig. 6c**).

**Figure 6.**
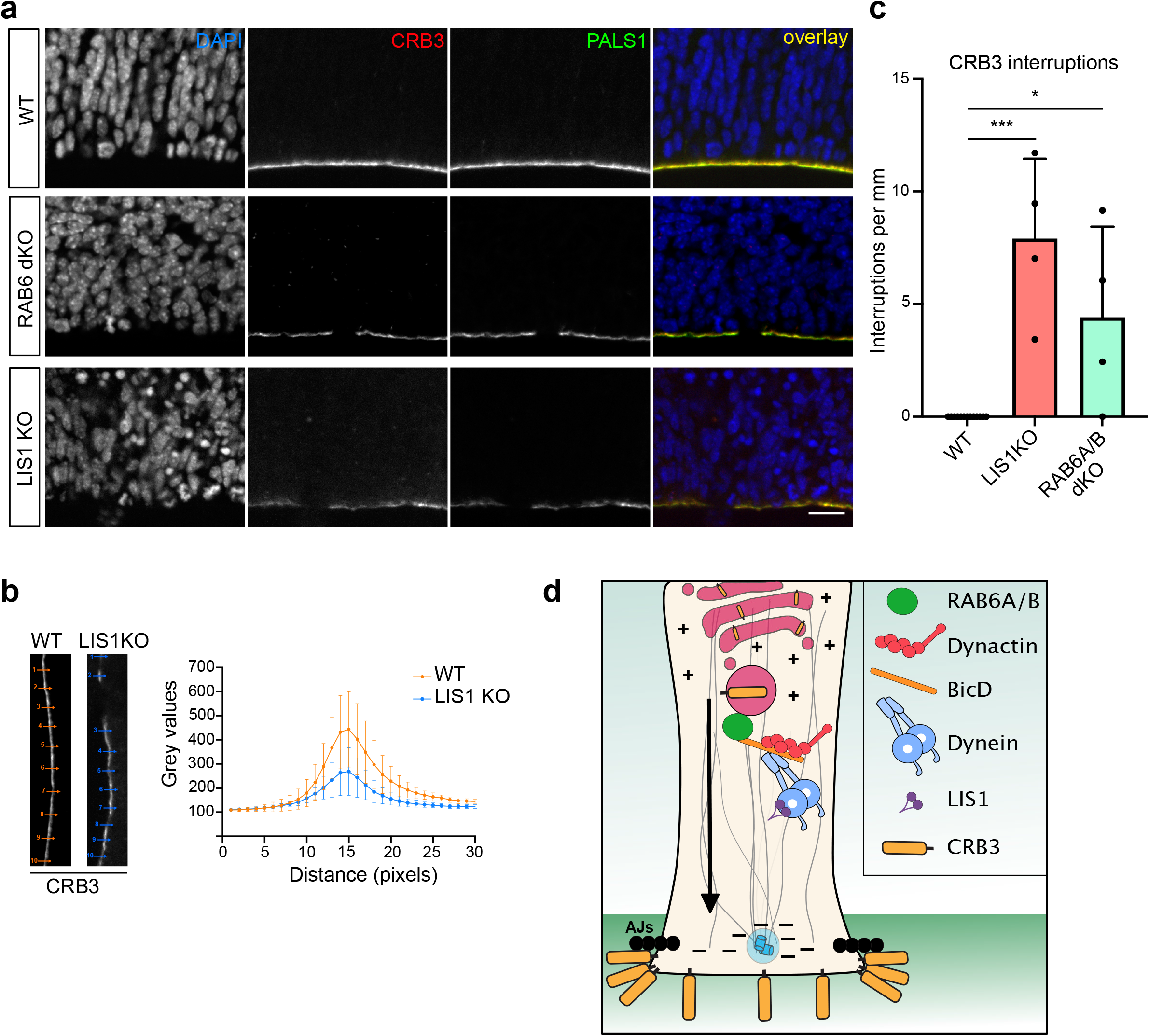
Apical localization of Crumbs in aRG cells depends on RAB6A/B and LIS1. **a.** CRB3 and PALS1 staining in WT, *RAB6A/B* dKO (E15.5) and *LIS1* KO (E12.5) brains. Scale bar = 25µm. **b.** CRB3 average apical signal intensity +/- SEM in WT and *LIS1* KO E12.5 brains. N= 3 brains. **c.** Quantification of CRB3 staining interruptions along the ventricular boundary of WT (E12.5 and E15.5; N=13 brains), *LIS*1 KO (E12.5, N=4 brains)) and *RAB6A/B* dKO (E15.5, N=4 brains). Kruskal-Wallis test with a Dunn *post-hoc* test and Benjamini-Hochberg procedure, *** p ≤ 0.001. **d.** Model for post-Golgi apical transport of CRB3 to the apical surface of aRG cells.

*RAB6A/B* dKO embryonic cortices also displayed an altered apical localization of CRB3 (**Fig. 6a**). As observed in *LIS1* KO brains, empty patches devoid of CRB3 and PALS1 occurred at a frequency of 4.3 per mm in *RAB6A/B* dKO (**Fig. 6c**). On the other hand, single gene depletion of *RAB6A* or *RAB6B* had no effect (data not shown). Altogether, the above results indicate that the RAB6-dynein-LIS1 apical trafficking pathway is required for proper transport of CRB3 and correct apical localization of the Crumbs complex.

## Discussion

The main finding of this study is that, in aRG cells, post-Golgi apical trafficking occurs in the microtubule minus end direction, via the RAB6-dynein-LIS1 complex, and is required for the apical localization of the Crumbs complex. As a consequence, genetic inactivation of *RAB6A/B* or *LIS1* leads to CRB3 loss at the ventricular surface and a delamination of aRG cells, which adopt features of bRG-like cells, including the ability to proliferate. We also establish aRG cells as a powerful epithelial model, enabling to resolve transport events in real time *in situ*.

### Post-Golgi transport is highly bidirectional in aRG cells

Dynein is largely viewed as a retrograde motor, driving trafficking towards the centre of the cell. We show here that in epithelial cells, where microtubule minus ends concentrate apically, dynein controls exit from the Golgi apparatus and transport to the apical surface. We observed that apical transport is however highly bidirectional, with RAB6+ vesicles constantly alternating in the apical and basal directions. Therefore, rather than being transported in a strictly polarized manner, RAB6+ vesicles actively oscillate, increasing the chances of reaching and docking to the apical surface. In non-polarized epithelial cells, although bi-directional movement can be observed, the trafficking of post-Golgi RAB6+ vesicles is largely unidirectional, moving towards the cell periphery in a kinesin-dependent manner^25, 50, 52^. The higher rate of minus end runs in aRG cells may point to a specific regulation of motors on RAB6+ vesicles upon epithelial polarization. Bicaudal family members, which recruit and activate dynein onto RAB6+ vesicles, are promising candidates for such regulation. Knock-out of *BICD2* in the mouse neocortex leads to ectopically dividing progenitors, phenocopying *LIS1* and *RAB6A/B* dKO, and suggesting apical polarity defects and delamination^53^. Transport in the minus end direction may be further biased by BICDR1, which is able to recruit 2 dynein molecules for faster movement, and induces strong accumulation of RAB6+ vesicles at the microtubule minus ends^31, 42, 54^.

### The RAB6-dynein-LIS1 complex controls post-Golgi apical transport of CRB3

Newly synthetized cargoes can traffic directly from the Golgi to the plasma membrane, though passage through intermediate recycling compartments was also proposed. We recently demonstrated that RAB6 acts as a general regulator of protein secretion and confirm here that CRB3 traffics within RAB6+ vesicles^23^. Because RAB6+ vesicles directly fuse with the plasma membrane, via its docking factor ELKS^50^, we favor a model whereby CRB3 is directly transported from the Golgi to the apical surface. CRB is known to be further maintained apically through a RAB11-dependent recycling route^55^. Retromer-dependent transport back to the TGN was also described, indicating that the RAB6-dynein-LIS1 pathway we describe here may also play a role in CRB recycling^56^. Of note, RAB6+ vesicles were also abundant in the basal process of aRG cells, but the mechanism(s) for sorting of apical and basal post-Golgi cargoes will require further investigation.

### RAB6A and RA6B redundantly control polarized trafficking

We observed that, unlike double KO, single deletion of *RAB6A* or *RAB6B* did not affect brain development, indicating that they were largely acting redundantly. Such redundancy was previously observed in cultured neurons following shRNA-mediated knockdown, as well as in MDCK cells where the very low levels are RAB6B are sufficient to compensate for *RAB6A* KO^24, 42, 54^. We also show that RAB6A/A’ and RAB6B act redundantly to control proper neuronal positioning, which may be caused by altered trafficking of adhesion molecules, including integrins^45^.

### Impaired apical post-Golgi trafficking leads to bRG-like cell production

bRG cells are generated from aRG cells and their amplification is a hallmark of gyrencephaly. The expression of several factors is known to affect their production but the underlying mechanisms remain largely unclear^57–59^. aRG cells were proposed to detach due to mitotic spindle rotation, or downregulation of the adherens junctions^15, 60, 61^. Recent evidence has demonstrated that delamination can be associated with Golgi structure abnormalities, and that detached aRG cells can reintegrate into the epithelium at early developmental stages but not at later neurogenic states^62, 63^. Here, using live imaging, we demonstrate that altered post-Golgi transport leads to a detachment of the apical process of aRG cells during interphase, and to the production of ectopically localized cells that maintain RG identity and proliferative capacity. These cells however appear to retract their basal process, likely due to impaired integrin-based contact at the basal end-foot. We did not observe the appearance of folding patterns in KO brains, due to the presence of an apoptosis-dependent microcephaly phenotype.

In conclusion, our results indicate that the maintenance of epithelial integrity during neocortex development relies on post-Golgi transport to the apical surface of aRG cells. This pathway can control the balance between aRG cell maintenance and bRG cell production, highlighting a potential site of action for factors that stimulate bRG cell production.

---

**Extended data Figure 1 (related to Figure 1).**
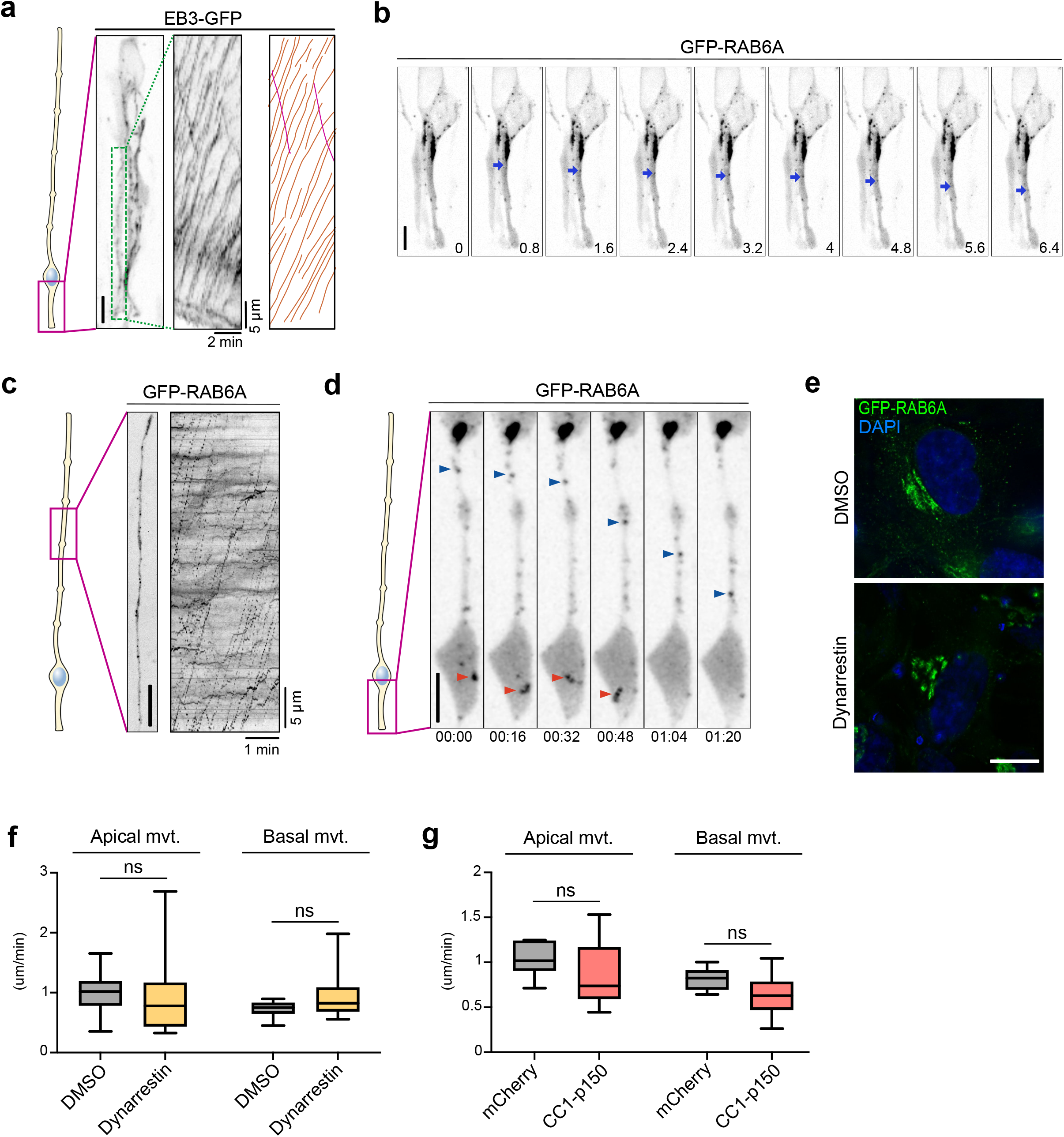
Microtubule polarity, RAB6A dynamics and dynarrestin validation. **a.** Live imaging of EB3-GFP in the apical process of an aRG cell at E15.5. Center: kymograph. Left: manual tracking of EB3 comets. Orange: basally-growing. Pink: Apically-growing. Scale bar = 5µm. **b.** Live imaging of GFP-RAB6A in the apical process of an aRG cell at E15.5. At 0.8 seconds a tubule is budding from the Golgi, leading to the formation of an apically-moving vesicle. Blue arrowhead indicates RAB6A+ vesicle. Scale bar = 5µm. **c.** Live imaging of GFP-RAB6A in the basal process of an aRG cell at E15.5. Right: kymograph. Scale bar = 5µm. **d.** Live imaging of GFP-RAB6A in the apical process of an aRG cell at E15.5. Red arrowhead: a RAB6A+ vesicle can be seen disappearing in the endfoot, suggesting fusion with the apical membrane. Blue arrowhead: a RAB6A+ vesicle moving apically within the apical process. Scale bar = 10µm. **e.** RPE-1 cells transfected with GFP-RAB6A to visualize the Golgi apparatus architecture, and treated for 4 hours with 100 µM dynarrestin or DMSO. Scale bar = 10µm. **f.** Velocity of apically and basally-moving RAB6A vesicles within the apical process of DMSO and dynarrestin-treated aRG cells. 142 vesicles from N=7 cells for DMSO, 74 vesicles from N=18 cells for dynarrestin. **g.** Velocity of apically and basally-moving RAB6A vesicles within the apical process of mcherry control and CC1-p150-expressing aRG cells. 120 vesicles from N=17 cells for mCherry control, 39 vesicles from N=11 cells for CC1-p150.

**Extended data Figure 2 (related to Figures 2 and 3).**
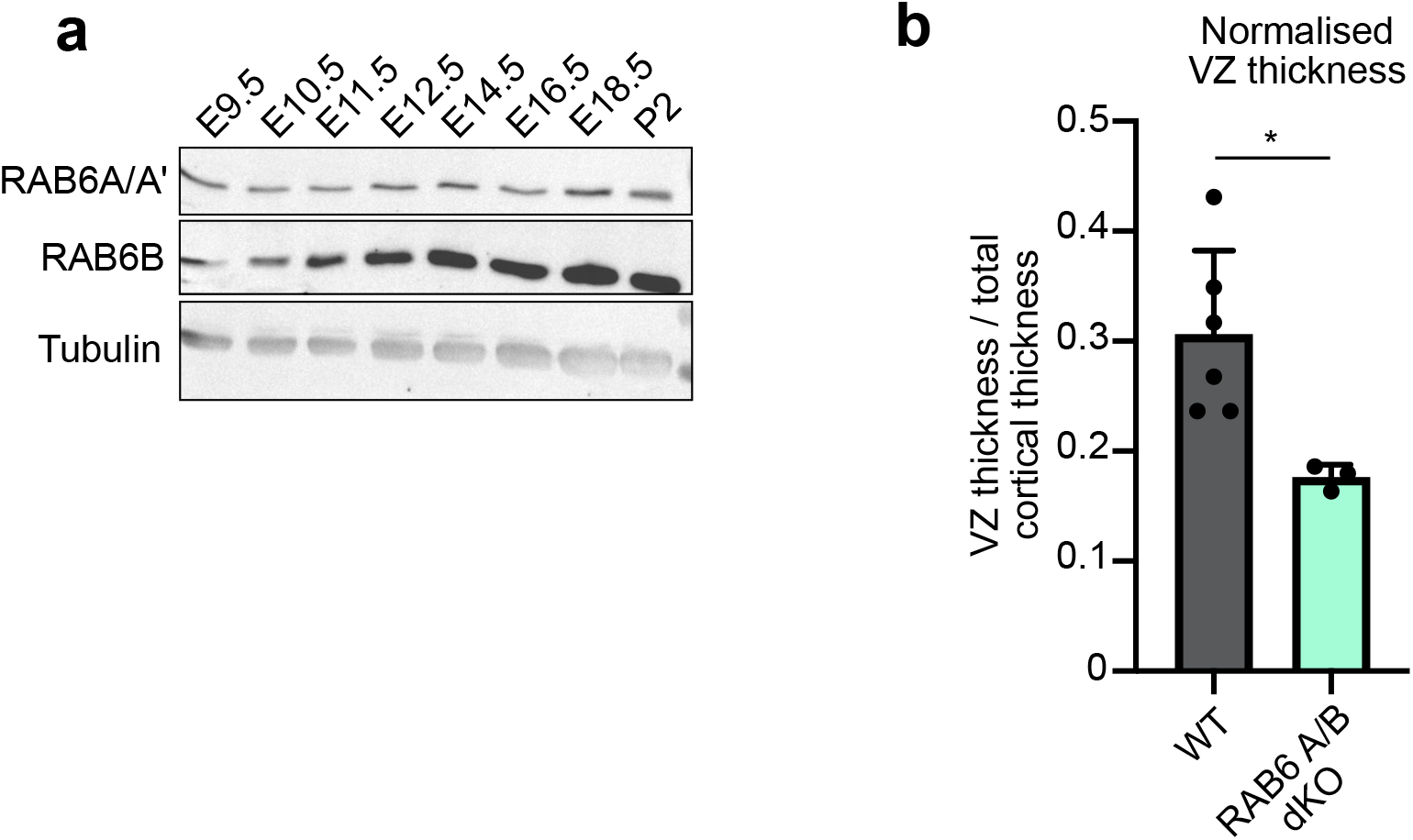
RAB6 expression in the brain and *RAB6A/B* dKO effect on VZ. **a.** RAB6A/A’ and RAB6B expression in the developing brain and at P2. **b.** Ventricular zone (VZ) thickness normalized to total cortical thickness in N=6 WT brains and N=3 *RAB6A/B* dKO. Mann–Whitney *U* test, * p≤0.05.

**Extended data Figure 3 (related to Figure 6).**
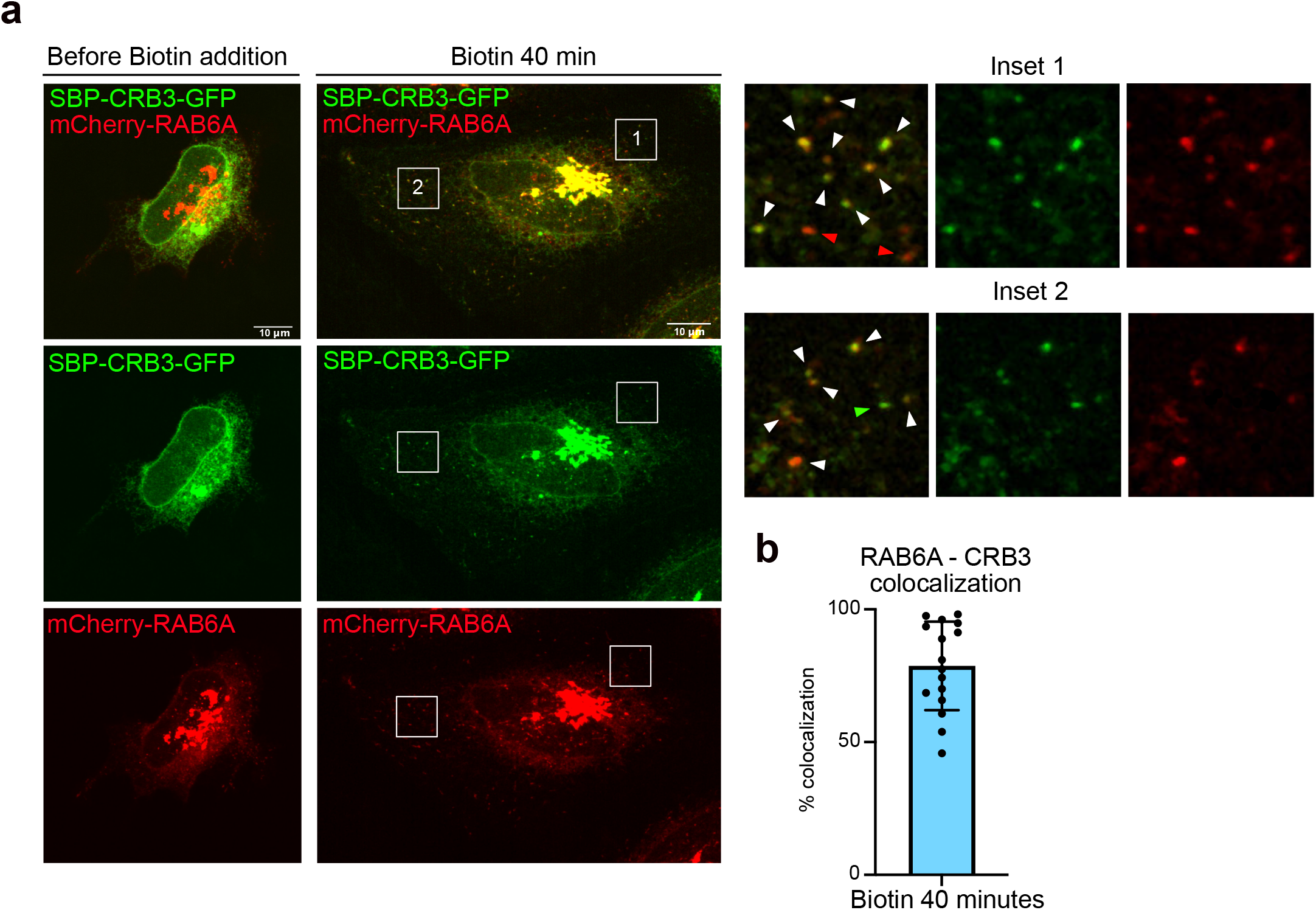
CRB3 exits the Golgi within RAB6+ vesicles. **a.** SBP-CRB3-GFP and mCherry-RAB6A localization in HeLa cells before and 40 minutes after addition of biotin. Right: Inset. White arrowheads: colocalizing foci. **b.** Quantification of SBP-CRB3-GFP and mCherry-RAB6A colocalization away from the Golgi apparatus 40 minutes after biotin addition.

**Supplemental Video 1 (related to Figure S1a). Live imaging of EB3-GFP in the apical process of an E15.5 aRG cell.** The vast majority of microtubules grows in the basal direction.

**Supplemental Video 2 (related to Figure S1b). Live imaging of GFP-RAB6A in the apical process of an E15.5 aRG cell.** A RAB6A+ vesicle can be seen budding from the Golgi and migrating apically (blue arrow).

**Supplemental Video 3 (related to Figure 1e). Live imaging of GFP-RAB6A in the apical process of an E15.5 aRG cell.** RAB6A+ vesicles can be seen moving bidirectionally.

**Supplemental Video 4 (related to Figure S1c). Live imaging of GFP-RAB6A in the basal process of an E15.5 aRG cell.** RAB6A+ vesicles largely move basally.

**Supplemental Video 5 (related to Figure S1d). Live imaging of GFP-RAB6A in the apical process of an E15.5 aRG cell.** A RAB6A+ vesicle disappears within the apical endfoot (red arrowdead). A RAB6A+ vesicle moves apically (blue arrowhead).

**Supplemental Video 6 (related to Figure 1i). Live imaging of GFP-RAB6A in the apical process of an E15.5 aRG cell.** RAB6A+ vesicles can be seen moving bidirectionally.

**Supplemental Video 7 (related to Figure 1i). Live imaging of GFP-RAB6A in the apical process of an E15.5 aRG cell treated with Dynarrestin.** The amount of RAB6A+ vesicles within the apical process is greatly reduced.

**Supplemental Video 8 (related to Figure 1i). Live imaging of GFP-RAB6A in the apical process of an E15.5 aRG cell expressing CC1-p150.** The amount of RAB6A+ vesicles within the apical process is greatly reduced.

Supplemental Video 9 (related to Figure 3g). Live imaging of an mCherry-expressing E17.5 aRG cell from a *RAB6A^loxP/loxP^; RAB6B^-/-^* genetic background (control). The apical process remains attached during the course of the movie.

Supplemental Video 10 (related to Figure 3g). Live imaging of an mCherry + CRE-expressing E17.5 aRG cell from a *RAB6A^loxP/loxP^; RAB6B^-/-^* genetic background (dKO). The apical process detaches during the course of the movie.

## Methods

### Animal breeding and care

All experiments involving mice were carried out according to the recommendations of the European Community (2010/63/UE). The animals were bred and cared for in the Specific Pathogen Free (SPF) animal facility of Institut Curie (agreement C75-05-18). All animal procedures were approved by the ethics committee of the Institut Curie (CEEA-IC #118) and by the French Ministry of Research (APAFiS# 26880-20200813165686-v1) in compliance with the international guidelines.

### Mice

#### Generation of RAB6B knockout mice

The constitutive *RAB6B* knock-out mice have been engineered using CRISPR/Cas9 technology to create a frame shift in the coding sequence. Two gRNA couples respectively targeting exons 2 and 3, and 2 and 4 were selected using the http://crispr.mit.edu/. gRNAs and Cas9m RNA were prepared according to the online protocol from Feng Zhang, https://www.addgene.org/crispr/zhang/. Briefly, the forward and the reverse oligonucleotides specific for the selected gRNA sequences were annealed and cloned into px330 plasmid. To get Cas9 mRNA and gRNAs, an *in vitro* transcription was performed on Cas9 pCR2.1-XL plasmid and gRNA plasmid using a T7 promoter, and the mMessage mMachine T7 ULTRA kit and MEGAshortscript T7 kit (Life Technologies), respectively. Cas9 mRNA and sgRNAs were then purified using the MEGAclear Kit (Thermo Fisher Scientific) and eluted in RNAse-free water. The gRNAs and Cas9mRNA quality were evaluated on agarose gel.

Eight-week-old superovulated B6D2F1/J (C57BL/6J × DBA/2J) females from Charles River France were superovulated by intraperitoneal (i.p.) administration of 5 IU of Pregnant Mare Serum Gonadotropin followed by an additional i.p. injection of 5 IU Human Chorion Gonadotropin 48 hours later. Superovulated females were mated to stud males of the same background. Zygotes were collected from the oviduct and were cultured in Cleave medium (Cook, K-RVCL-50) at 37°C under 5% CO2 until microinjection. An injection solution was prepared as following: Cas9 mRNA at 100 ng/µl and 50ng/µl for each gRNA in Brinster buffer (10 mM Tris-HCl pH 7.5; 0.25 mM EDTA) and passed through 0.22 µm pore size filter. Cytoplasmic microinjection was performed into mouse fertilized oocytes. Microinjected embryos were transferred into 0.5 dpc NMRI pseudo-pregnant females with 12 zygotes per oviduct. Selected founders F0 carrying a 1 bp deletion in exon 2 and a 279 bp inversion, both leading to a premature STOP codon, were then backcrossed to C57BL6/N to segregate out undesired genetic events.

#### RAB6A/B dKO and LIS1 KO

*RAB6A^loxP/loxP^* mutant mice were previously generated and characterized^46^. *RAB6A^loxP/loxP^* mice were first crossed with *RAB6B^-/-^* mice to generate *RAB6A^loxP/loxP^*; *RAB6B^-/-^* animals, which were viable and fertile. These animals were then crossed with *Emx1-Cre* (JAX 005628) animals to generate *Emx1*; *RAB6A^loxP/loxP^*; *RAB6B^-/-^* (*RAB6A/B* double knockout) animals. *LIS1* conditional knockout mice (*LIS1^-^*^/-^, also known as Pafah1b1-loxP^64^) were crossed with *Emx1-Cre* mice.

### *In utero* electroporation of mouse embryonic cortex

Pregnant mice were anesthetized with isoflurane gas, and injected subcutaneously first with buprenorphine (0.075 mg/kg) and a local analgesic, bupivacaine (2 mg/kg), at the site of the incision. Lacrinorm gel was applied to the eyes to prevent dryness/irritation during surgery. The abdomen was shaved and disinfected with ethanol and antibiotic swabs, then opened, and the uterine horns exposed. Plasmid DNA mixtures were used at a final concentration of 1 µg/µl per plasmid (except for GFP-RAB6 overexpression experiments: mCherry empty vector/ CC1-p150 1.5 µg/µl versus GFP-RAB6A 250 ng/µl), dyed with Fast Green and injected into the left lateral ventricle of several embryos. The embryos were then electroporated through the uterine walls with a NEPA21 Electroporator (Nepagene) and a platinum plated electrode (5 pulses of 50 V for 50 ms at 1 second intervals). The uterus was replaced and the abdomen sutured. The mother was allowed to recover from surgery and supplied with painkillers in drinking water post-surgery.

### Immunostaining of brain slices

Mouse embryonic brains were dissected out of the skull, fixed in 4% PFA for 1 hour, and 80 µm-thick slices were prepared with a Leica VT1200S vibratome in PBS. Slices were boiled in citrate sodium buffer (10mM, pH6) for 20 minutes and cooled down at room temperature (antigen retrieval). Slices were then blocked in PBS-Triton X100 0.3%-donkey serum 2% at room temperature for 1 hour, incubated with primary antibody overnight at 4°C in blocking solution, washed in PBS-Tween 0.05%, and incubated with secondary antibody for 2 hours at 4°C in blocking solution before final wash and mounting in aquapolymount. Imaging was performed on a spinning disk wide microscope equipped with a Yokogawa CSU-W1 scanner unit equipped with a with a 40X Apo-Plan objective. Whole brains were imaged using a Leica MZ8 Stereozoom Microscope.

### Western blots

Tissue extracts were performed from whole embryos (E9.5), heads (E10.5; E11.5; E12.5), brains (E14.5); or cortices (E15.5; E16.5; E18.5; P2) in 150mM NaCl, 50mM Tris pH8, 0,1% v/v SDS, 0,5% v/v Nonidet P40, 1X complete protease inhibitors, briefly sonicated, and centrifuged. Protein concentrations were measured using a BCA protein assay kit. 25mg were analysed in 10% acrylamide-containing gels.

### Expression constructs and antibodies

The following plasmids were used in this study: ManII-GFP, CC1-p150, Streptavidin-KDEL SBP-CRB3A-GFP (Franck Perez); GFP-RAB6A^52^; EB3-GFP (gift from Matthieu Piel); mCherry2-C1 vector (gift from Michael Davidson, Addgene plasmid #54563); Cre (gift from David Liu, Addgene plasmid #123133); pCAG-Cre-IRES2-GFP vector (gift from Anjen Chenn, Addgene plasmid #26646); pCAG-GFP vector (gift from Richard Vallee, Columbia University).

Antibodies used in this study were mouse anti-*γ*Tubulin (Sigma-Aldrich, T5326), rat anti-Crumbs3 (gift from André Le Bivic, Marseille), rabbit anti-MPP5/PALS1 (Proteintech, 17710-1-AP), human anti-GFP (recombinant antibody platform (Tab-IP) - Institut Curie, A-R-H#11), rabbit anti-Pax6 (Biolegend, B214847), goat anti-phospho-Histone 3 (Santa Cruz, SC-12927), mouse anti-phospho-Vimentin (Abcam, 22651), CUX-1 (Santa-Cruz, discontinued), rabbit anti-cleaved-Caspase 3 (Cell Signaling, 9661S), rabbit anti-RAB6A/A’ (home-made^65^), rabbit anti-RAB6B (Proteintech, 10340-1-AP), human anti-*α*Tubulin (recombinant antibody platform (Tab-IP) - Institut Curie, A-R-H#02). Secondary antibodies: donkey Alexa Fluor 488 anti-mouse, anti-rabbit, anti-goat (Jackson laboratories 715-545-150, 711-165-152, 715-605-152), donkey Alexa Fluor 555 anti-mouse, anti-rabbit, anti-goat (Jackson laboratories 715-545-150, 711-165-152, 715-605-152), donkey Alexa Fluor 647 anti-mouse, anti-rabbit, anti-goat (Jackson laboratories 715-545-150, 711-165-152, 715-605-152).

### Subcellular live imaging in mouse embryonic brain cortex slices

To record GFP-RAB6A dynamics in radial glia *in situ*, we used the following approach. 24 hours after the electroporation of E15.5 to E16.5 embryos, the pregnant mouse was sacrificed and the electroporated embryos recovered. Brains were dissected in artificial cerebrospinal fluid (ACSF) and 250 µm-thick coronal slices were prepared with a Leica VT1200S vibratome in ice-cold ACSF. The slices were cultured on membrane filters over enriched medium (DMEM-F12 containing B27, N2, 10 ng/ml FGF, 10 ng/ml EGF, 5% fetal bovine serum and 5% horse serum). After recovery in an incubator at 37°C, 5% CO2 for 2 hours (or 48 hours for human tissue to allow for construct expression), the filters were cut and carefully turned over on a 35 mm FluoroDish (WPI), in order to position the sample in direct contact with the glass, underneath the filter (to maintain the sample flat).

Live imaging was performed on a fully motorized spinning disk wide microscope driven by Metamorph software (Molecular Devices) and equipped with a Yokogawa CSU-W1 scanner unit to increase the field of view and improve the resolution deep in the sample. The inverted microscope (Nikon Eclipse Ti2) was equipped with a high working distance (WD 0.3 mm) 100X SR HP Plan Apo 1.35 NA Silicon immersion (Nikon) and a Prime95B sCMOS camera (Photometrics). To maintain stable cell culture conditions (37°C, humidity, 5% CO_2_), time-lapse imaging was performed on a STX stage top incubator (Tokai Hit). Z-stacks of 3-5 µm range were taken on a Mad City Lab piezo stage (Nano Z500) with an interval of 1 µm. Maximum projections were generated from which kymographs were generated. Tracking and quantifications of GFP-RAB6A+ vesicle dynamics were directly performed on the movies and the kymographs were used for validation and display purposes. Videos were mounted in Metamorph. Kymograph generation (KymographBuilder), Tracking of GFP-RAB6A+ vesicles (manual tracking) as well as image modifications (brightness and contrast, background, gamma) were carried out on Fiji. Figures were assembled in Affinity Designer.

### RUSH assay *in situ*

E15.5 to E16.5 embryos were electroporated with a Streptavidin-KDEL SBP-CRB3-GFP construct with or without CC1-p150 for 24 hours. Slicing and culture were performed as for subcellular live imaging experiments. Biotin was added to the enriched medium (40µM final) for the indicated period of time (37°C, 5% CO2) prior to PFA fixation. Immunostaining against GFP was performed to amplify fluorescence (see immunostaining section) prior to mounting.

### Statistical analysis

All the statistical analysis has been made using R 4.0.5. R Core Team (2021), R Foundation for Statistical Computing, Vienna, Austria (https://www.R-project.org/). Due to the low sample sizes inherent to *in vivo* work, we conducted non-parametric analyses. Median comparisons between 2 conditions have been made with a Mann–Whitney *U* test (Fig. 1j-m, 3b, 3d, 4c, 4d and S2b). When more than 2 conditions were compared, we used Kruskal-Wallis test with a Dunn *post-hoc* test and Benjamini-Hochberg procedure to control the false discovery rate using the dunn.test package (Fig. 2c, 2e, 3e and 6c). These analyses have been made considering the animal as the statistical unit except for the figures 1j-1m. Embryos for a given condition come from different litters. For categorical data (Fig. 3h, 5d-e) and data from figures 1j-1m, S1f-g, we considered each cell as a statistical unit. Since the cells are electroporated *in-situ*, we made the reasonable approximation that cells received their constructs independently and their properties are measured individually at the cell scale. We validated this hypothesis by repeating experiments in different independent animals to conclude that the effect was not due to cells coming from biased individuals due to an abnormal electroporation or an abnormal embryo. For categorial data, analysis has been made using a Fisher’s exact test (Fig. 3h) accompanied with a Benjamini-Hochberg procedure to control the false discovery rate when more than 2 conditions were compared (Fig. 5d-e). These categorical data are depicted as percentages for clarity. *p*-values superior to 0.05 are considered as not significant.

## Acknowledgements

The authors greatly acknowledge the Cell and Tissue Imaging (PICT-IBiSA), Institut Curie, member of the French National Research Infrastructure France-BioImaging (ANR10-INBS-04) and the Nikon BioImaging Center (Institut Curie, France). We greatly acknowledge the Recombinant Antibody Platform of the Institut Curie for the production of antibodies. We thank D. Massey-Harroche and A. Le Bivic for the anti-Crumbs3 antibody. A.D.B is an INSERM researcher. This work was supported by the ANR (ANR-19-CE13-0002-02), CNRS, Institut Curie, the Ville de Paris “Emergences” program, Labex CelTisPhyBio (11-LBX-0038), and PSL.

## Author contributions

J.B.B, B.G and A.D.B conceived the project. J.B.B and A.D.B analyzed the data. J.B.B, B.G and A.D.B wrote the manuscript. F.E.M generated the *RAB6B* KO mouse lines. J.A.C and L.C sequenced the *RAB6B* KO mouse lines and assisted with surgery and *in utero* electroporation. S. Baloul performed RUSH assays *in situ*. S. Bardin supervised the RAB6 mouse mutant colonies, performed all the crossings to generate *RAB6A/B* dKO animals and performed the western-blots. M.P generated the *LIS1* KO embryos. M.L and S.M.L performed and quantified the RUSH assay in HeLa cells. G.B and F.P designed and supervised the RUSH assays. V.S assisted with high resolution *in situ* imaging. H.L performed all statistical analyses.

## Notes

### Competing Interest Statement

The authors have declared no competing interest.

